# Probiotic-conditioned microbiota from preterm infants modulate immune response to pathogen challenge in a humanized mouse model

**DOI:** 10.64898/2025.12.08.692141

**Authors:** Justine Smout, Till-Robin Lesker, Lisa Hoenicke, Diego Ortiz, Mangge Zou, Friederike Kruse, Sabine Pirr, Maike Willers, Christoph Härtel, Christine Falk, Natalia Torow, Dorothee Viemann, Till Strowig, Jochen Huehn

## Abstract

Early-life host-microbe interactions critically shape immune development, lifelong homeostasis, and disease susceptibility. The PRIMAL trial (Priming Immunity at the Beginning of Life) demonstrated that multistrain probiotics shifted the gut microbiota of very preterm infants toward eubiosis without affecting sepsis incidence, yet the immunological consequences remained unresolved. To explore this, we colonized germ-free female mice with fecal samples from probiotic- or placebo-treated preterm infants from the PRIMAL trial. Microbiota composition was vertically transmitted and stable across generations. At steady-state, 3-week-old pups colonized with probiotic-conditioned microbiota exhibited markedly reduced populations of innate immune cells, particularly in the colon, with subtler effects in the small intestine and spleen, while adaptive immune subsets were less affected. Upon enteropathogenic *Escherichia coli* challenge at day 5, pups harboring probiotic-conditioned microbiota displayed reduced growth and impaired bacterial clearance, correlating with diminished numbers of key innate immune cell populations. These findings demonstrate that probiotic-driven shifts in human-derived microbial communities can attenuate immune cell development in mice and alter early-life infection outcomes. Our study underscores the complex, context-dependent effects of probiotics on the neonatal microbiota–immune axis and provides mechanistic insight into how interventions in preterm infants may influence susceptibility to infection.

## Introduction

The acquisition and development of the gut microbiota during early life follow successive waves of exposure to microorganisms, which play a vital role in establishing the interactions between hosts and microbes that are essential for optimal symbiosis and future health (1, 2). Although substantial interindividual variation is observed during this initial colonization phase, the early-life microbiota is generally characterized by low diversity and density, rendering it highly susceptible to exogenous perturbations (3). Moreover, the initial absence of competitors for space and nutrients facilitates a rapid increase in bacterial numbers in the gut (4, 5). The most significant microbial inoculum occurs at birth and shortly thereafter, through the opportunistic colonization of the first bacterial species encountered by an infant in its environment, and also via vertical transmission from mother to infant (4, 6, 7). This early microbial establishment shapes the trajectory of the gut microbiota and plays a fundamental role in immune system development.

Over the past decade, studies have highlighted the crucial role of early microbial encounters in shaping the host’s immune system, metabolism, and resistance to pathogen colonization (8, 9). These interactions are vital throughout life but are especially important during the neonatal period, which coincides with a limited window of microbial permissiveness and immune training known as the “window of opportunity.” During this time, the initial colonizers, including both commensals and pathogens, have a particularly strong impact on the immune system (5–7, 10).

Gestational age is one of several factors that influence the composition of the gut microbiota (1). Preterm infants are anatomically and immunologically immature, exhibiting a delay in the acquisition and development of their gut microbiome (11). Compared with term infants, they show reduced microbial diversity, delayed colonization by beneficial anaerobes, and an early dominance of facultative and potentially pathogenic taxa, largely shaped by antibiotic exposure, limited maternal microbial transfer, and the NICU environment (12, 13). This atypical microbial assembly coincides with an underdeveloped mucosal and systemic immune system (11, 14, 15). Together, this can result in altered host-microbiome interactions that delay postnatal immune maturation, increasing the susceptibility of preterm infants to severe infections such as late-onset sepsis and necrotizing enterocolitis (16–18). Several studies have shown that in preterm infants initial gut colonizers can even be the primary source of the invading pathogens causing sepsis, particularly gram-negative pathobionts such as *Escherichia coli* (19, 20). This underscores the clinical relevance of the interplay between the preterm gut microbiome and preterm gut barrier function, which if heavily impaired results in severe gut-derived systemic infections.

Studies in humans have associated specific microbial species with reduced incidence and improved outcomes of infection in preterm infants, highlighting the potential of probiotics for prophylactic and adjunctive therapeutic interventions (21–23). Recently, the multicenter phase 3 *Priming Immunity at the Beginning of Life* (PRIMAL) randomized clinical trial (RCT) assessed the impact of probiotic intervention in preterm infants to prevent colonization by resistant bacteria and to shape their microbiome. In this study, 618 infants (28–32 weeks gestation) received a 28-days multistrain probiotic or placebo. (24, 25). While the intervention did not reduce colonization by multidrug-resistant organisms by day 30, it significantly shifted the gut microbiota toward a more eubiotic composition resembling that of term infants (24). However, detailed insights into how age-related programming and microbiota-dependent imprinting influence immune maturation and long-term susceptibility to infectious diseases are still lacking.

To gain deeper insight into the effects of early-life probiotic supplementation independently of the direct actions of probiotic strains, we established a fecal microbiota transplantation (FMT) model in germ-free mice using fecal samples from preterm infants who had received either probiotics or placebo during the first month of life. Importantly, the mice were never exposed to probiotics themselves; instead, they were colonized with microbiota that had been previously shaped by probiotic supplementation. This design allowed us to investigate how probiotic-conditioned differences in the neonatal microbiota influence immune development in early life. We studied the immune system of these ‘humanized’ pups under homeostasis and then subjected them to an enteropathogenic *Escherichia coli* (EPEC) challenge in order to evaluate the impact of the microbiota on early-life infection. This approach has enabled us to develop a preterm-derived humanized mouse model that provides a tool to examine how early-life microbial ecosystems shape immune development and infection outcomes.

## Materials and Methods

### Ethics statement

All animal experiments have been performed in agreement with the guidelines of the Helmholtz Centre for Infection Research, Braunschweig, Germany, the National Animal Protection Law (Tierschutzgesetz) and Animal Experiment Regulations (Tierschutz-Versuchstierordnung), and the recommendations of the Federation of European Laboratory Animal Science Association (FELASA). The study was approved by the Lower Saxony State Office of Nature, Environment and Consumer Protection (LAVES), Oldenburg, Germany; permit number 33.8-42502-04-22-00247.

### Mice

Germ-free (GF) C57BL/6NTac mice were bred in isolators (Getinge) at the germ-free facility of the Helmholtz Centre for Infection Research (HZI, Braunschweig, Germany). Sterilized food and water were provided *ad libitum*, and the mice were maintained on a strict 12-hour light/dark cycle. Euthanasia was performed by CO2 asphyxiation for the 3-week-old and adult mice and by decapitation after prior anesthesia with inhaled isoflurane for the postnatal day 12 mice.

### Human samples

Preterm stool samples (≤32 weeks of gestational age) were collected at day of life 30 as part of the PRIMAL study conducted at the Medical School Hannover (25). Briefly, participants were randomized within the first 48 hours after birth to receive either a multi-strain probiotic or a placebo in a 1:1 ratio. Probiotic or placebo administration continued for 28 days. The probiotic consisted of *Bifidobacterium animalis* subsp. *lactis* (BB-12), *Bifidobacterium longum* subsp. *infantis*, and *Lactobacillus acidophilus* (La-5), provided in daily dose capsules containing 1.5 × 10^9^ colony-forming units (CFUs) of each strain per capsule. The placebo consisted of cornstarch powder encapsulated to match the appearance and odor of the probiotic formulation.

For this study, stool samples were solubilized, aliquoted, and stored at -80°C in anaerobic brain heart infusion (BHI) medium containing 50% glycerol (Carl Roth) and palladium black (Sigma-Aldrich). The solubilized samples were cultured on BHI agar plates (Oxoid and BD Bioscience) supplemented with 5% sheep blood (Thermo Fisher Scientific) and 0.1% vitamin K3 (Carl Roth). Both aerobic and anaerobic incubation conditions were employed to quantify the CFUs of culturable bacteria per gram of feces.

### Human neonatal microbiota transplantation

Six- to 8-week-old GF female mice were colonized via oral gavage with 5 × 10^7^ CFU/mL of preterm microbiota feces diluted in sterile PBS (200 µL per mouse). Two weeks after microbiota transplantation, the females were mated with GF males. Once pregnancy was confirmed (approximately two weeks later), each female was housed in an individual cage. Vertical transmission of the human microbiota from the parental generation (mothers) to their offspring (F1 generation) was monitored. Microbiota engraftment in both generations was assessed at the time of sampling using 16S rRNA sequencing.

### Neonatal infection model with enteropathogenic *Escherichia coli*

Wild-type enteropathogenic *Escherichia coli* (EPEC) strain E2348/69 (Streptomycin^R^) was inoculated and plated on Luria Broth (LB) agar supplemented with 100 µg/mL streptomycin (Sigma-Aldrich).

Single colonies were picked and cultured for 8 hours at 37°C with shaking at 200 rpm. Four- to 5-day-old F1 generation mice were then infected via intragastric injection with 1 × 10^7^ CFU, diluted in 50 µL of sterile PBS. The infection course was monitored daily, and the animals were euthanized 7 days post-infection. Pathogen burden was assessed by plating homogenized organs and intestinal contents on LB agar containing 100 µg/mL streptomycin to selectively culture EPEC. CFUs were quantified and expressed as CFU/organ. Spleen and small intestine (SI) were collected for flow cytometry analysis.

### 16S rRNA analysis of microbial communities

Fresh stool samples or colonic contents were collected and immediately stored at −80°C until processing. DNA was isolated following an established protocol (26). Briefly, 500 μL of extraction buffer (200 mM Tris, 20 mM EDTA, 200 mM NaCl, pH 8.0) (Roth), 200 μL of 20% SDS (AppliChem), 500 μL of phenol:chloroform:isoamyl alcohol (PCI) (24:24:1) (Roth) and 100 μL of zirconia/silica beads (0.1 mm diameter) (Roth) were added to each sample. Bacterial lysis was achieved via mechanical disruption using a Mini-BeadBeater-96 (BioSpec) for two 2-minute cycles. The aqueous phase was recovered after centrifugation and subjected to an additional PCI extraction. DNA was then precipitated by adding 500 µL of isopropanol (J.T. Baker) and 0.1 volume of 3 M sodium acetate (AppliChem), followed by incubation at −20°C for several hours or overnight. Precipitated DNA was pelleted by centrifugation at maximum speed (4°C, 20 minutes), washed, and dried using a vacuum centrifuge. The DNA pellet was resuspended in TE buffer (AppliChem) containing 100 µg/mL RNase I (AppliChem). To remove PCR inhibitors, crude DNA was further purified using a column purification kit (BioBasic Inc.). 16S rRNA gene amplification targeting the V4 region (F515/R806) was performed following the protocol established by Caporaso et al. (27). DNA was normalized to 25 ng/µL and used for PCR with unique 12-base Golay barcodes incorporated via specific primers (Sigma). PCR was conducted in triplicate for each sample using Q5 polymerase (New England Biolabs) under the following conditions: initial denaturation at 98°C for 30 seconds, followed by 25 cycles of denaturation at 98°C for 10 seconds, annealing at 55°C for 20 seconds, and extension at 72°C for 20 seconds. PCR amplicons were pooled, normalized to 10 nM, and sequenced on an Illumina MiSeq platform using 250 bp paired-end sequencing (PE250). Sequencing reads were assembled, filtered, and clustered with the USEARCH v8.1 software package (http://www.drive5.com/usearch/). Low-quality reads were removed, and sample-specific barcodes were used to bin reads with QIIME v1.8.0 (28). Paired-end merging was performed with the fastq_mergepairs command (fastq_maxdiffs 30), and quality filtering was conducted using fastq_filter (fastq_maxee 1), with a minimum read length of 250 bp and a minimum of 1,000 reads per sample. Reads were clustered into operational taxonomic units (OTUs) at 97% identity through open-reference OTU picking, with representative sequences identified using the UPARSE algorithm (29). OTUs with abundance >0.5% were retained for analysis, and taxonomic classification was performed with the RDP Classifier at an 80% bootstrap confidence cutoff using the Greengenes or SILVA databases. Sequences lacking a reference match were assembled *de novo* with UCLUST. Phylogenetic relationships among OTUs were inferred using FastTree based on PyNAST-aligned sequences (30). The resulting OTU abundance table and mapping file were analyzed and visualized in the R statistical environment using the phyloseq package (31).

### Isolation of immune cells from different tissues

Spleens were collected and mechanically disaggregated through a 30 µm cell strainer. Splenocytes were subjected to erythrocyte lysis using a buffer containing 0.01 M potassium bicarbonate, 0.155 M ammonium chloride, and 0.1 mM EDTA (pH 7.5). The resulting cells were resuspended in PBS supplemented with 0.2% BSA and kept on ice until further processing.

Colons were collected, opened longitudinally, and washed with pre-digestion medium (PBS, containing 0.2% BSA (Merck) and 5 mM EDTA (Sigma-Aldrich)) to remove fecal content. Tissues were incubated at 37°C with continuous shaking (250 rpm) for 20 minutes. After predigestion, colons were cut into 1–2 mm pieces and digested in RPMI medium containing 1 mg/mL Collagenase D (Roche) and 0.1 mg/mL DNase I (Roche) at 37°C for 1 hour. SI were harvested, opened longitudinally and washed with PBS to remove feces. Peyer’s patches were not removed prior to processing. The lamina propria was dissociated using the mouse lamina propria dissociation kit (Miltenyi Biotec) with the gentleMACS™ Octo Dissociator and heaters, following the manufacturer’s instructions.

Colonic and intestinal cells were purified using a 40%/80% Percoll gradient (GE Healthcare) centrifugation at 780 × g for 20 minutes at room temperature, with acceleration and brake settings turned off. Cells from the interphase layer were collected, washed, resuspended in PBS containing 0.2% BSA, and kept on ice until further use.

### Flow cytometry

Prior to staining, cells were stimulated with phorbol 12-myristate 13-acetate (PMA, 10 ng/ml), Ionomycin (0.5 μg/ml), and Brefeldin A (10 μg/ml) (all Sigma-Aldrich) for 4 h at 37°C. Flow cytometric staining was performed as described recently (32). Briefly, single-cell suspensions were washed with PBS, and dead cells were labeled using a UV-excitable fixable Blue Dead Cell Stain kit (Thermo Fisher) for 15 minutes at 4°C. Cells were fixed and permeabilized using the Foxp3 Transcription Factor Staining Kit (Thermo Fisher) for 30 minutes at room temperature in the dark, followed by overnight intracellular staining at 4°C. The following antibodies were used: anti-CD3 (145-2C11, BV480), anti-CD11b (M1/70, BUV496), anti-CD45 (30-F11, BV786), anti-CD49a (Ha31/8, BUV615), anti-CD64 (X54-5/7.1, BV650), anti-CD103 (M290, BUV395), anti-CD127 (A7R34, RB780), anti-Gata3 (L50-823, AF647), anti-IgD (11-26c2.a, BV711), anti-IgM (II/41, BUV563), anti-IL-17A (TC11-1810.1, BV605), anti-Ly6G (1A8, BV750), anti-Rorγt (Q31-378, BV421), anti-Siglec F (E50-2440, APC-R700), and anti-Siglec H (440c, BUV661) from BD Biosciences; anti-CD4 (RM4-5, BV570), anti-CD8 (53-6.7, Spark Blue 550), anti-CD11c (N418, Pacific Blue), anti-CD19 (6D5, PE-Cy5), anti-γδTCR (GL3, APC-Fire750), anti-IL-10 (JES5-16E3, APC), anti-Ly6C (HK14, PE-Fire810), anti-MHCII (M5/114.15.2, BV510), anti-NK1.1 (PK136, PE-Dazzle594), anti-T-bet (4b10, PE-Cy7), and anti-XCR1 (ZET, PerCP-Cy5.5) from BioLegend; anti-CD45R (RA3-6B2, BUV737), anti-TCRβ (H57-597, PE-Cy5.5), anti-Foxp3 (FJK-16S, AF488), anti-IFN-γ (XMG1.2, PE), and anti-Mertk (DS5MMER, BUV805) from Thermo Fisher. Unconjugated anti-CD16/CD32 (BE0307) for blocking unspecific binding was purchased from BioXCell. Samples were acquired on a BD FACS Symphony A5 SE Cell Analyzer (5 lasers, 50 parameters, 48 fluorescence channels), and data were analyzed using FlowJo software (v10.10; FlowJo, LLC).

### Quantification of immunoglobulins in the cecum content

Fresh cecum contents were collected and immediately stored at −80°C until further processing. The concentration of mouse immunoglobulins isotypes IgG1, IgG2a, and IgG2b, IgG3, IgM, and IgA in the caeca were determined with the kit LEGENDplex™ Mouse Immunoglobulin Isotyping Panel (6-plex), according to the manufacturer’s instructions (BioLegend). Data acquisition was performed on a BD FACS Fortessa.

### Cytokine quantification in the serum

Blood samples were collected and centrifuged twice at 5,000 rpm for 10 minutes to obtain serum, and immediately stored at −80°C until further analysis. Cytokines and chemokines levels (IL-1α, IL-1β, IL-2, IL-3, IL-4, IL-5, IL-6, IL-9, IL-10, IL-12p40, IL-12p70, IL-13, IL-17A, Eotaxin, G-CSF, GM-CSF, IFN-γ, KC, MCP-1, MIP-1α, MIP-1β, RANTES, and TNF-α) were measured with the Bio-Plex Pro Mouse Cytokine 23-plex Assay (Bio-Rad) following manufacturer’s instruction.

### Statistical analysis

Graphs were generated using GraphPad Prism 10 (GraphPad Software; San Diego, USA) and R 4.3.1. Data are presented as mean ± SD. Statistical comparisons were performed using GraphPad Prism 10. Normality of the data was assessed to determine the appropriate statistical method. Statistical analysis was performed using an unpaired t test or Mann-Whitney test for 2-group comparisons, or an ordinary one-way ANOVA (Holm-Sidak’s multiple comparisons test) or Kruskal-Wallis test (Dunn’s multiple comparisons test) for more than 2 multiple comparisons. P-values ≤ 0.05 were considered statistically significant.

## Results

### Establishment of a neonatal humanized model for preterm microbiota engraftment

Because the acquisition of early life microbiota has been shown to be primarily due to maternal source communities (4, 33), we established a humanized neonatal mouse model of preterm infant microbiota using a vertical transfer system from adult mice to their pups. In brief, fecal samples from six preterm infants (**Table 1**) (three fecal samples from placebo-treated preterm infants and three fecal samples from probiotic-treated preterm infants) were orally administered to 6-week-old GF mice. Two weeks later, the colonized mice were mated, and their offspring were sampled 3 weeks after birth (F1 generation) as well as their respective mothers (parental generation) (**Figure 1A**). To validate our humanized mouse model, colon contents were collected, and the microbiota diversity and composition were evaluated using 16S rRNA amplicon sequencing. In terms of α-diversity, we observed similar bacterial richness in both parental and F1 generations among the 6 different donors, confirming the efficacy of vertical transfer from mother to offspring (**Figure 1B**). However, the richness between the humanized mouse samples and the corresponding human donor sample varied depending on the donor. While for donors 1 and 14, the colonized mice showed a higher richness compared to their human donor sample, the opposite was true for the other donors. Shannon diversity showed similar differences between the input, parental, and F1 generations (**Figure 1B**). In case of donor 1, the microbiome engraftment resulted in a similar pattern with increased Shannon diversity in both the parental and F1 generations compared to their human counterpart. However, when engrafted with microbiota originating from donor 2, the recipient mice showed a large reduction in diversity. These results suggest that differences in the engrafted microbiota compared to the original input are not only related to the presence and absence of OTUs but also to the dominance and expansion of specific bacteria in the mouse colonic niche (**Figure S1A**). Analyses based on the relative abundance of OTUs confirmed differences in microbial dominance between the human donors, parents, and F1 mice (**Figure 1C and S1B**). The genera *Blautia* and *Enterocloster* exhibited significant expansion and persistent domination of the murine colonic niche. In contrast, the relative abundances of other genera, particularly *Bifidobacterium* and *Enterococcus*, were found to be reduced in the mouse colon. An exception was observed for donor 11, for which the microbiota composition of recipient mice exhibited a high degree of similarity to that of the human donors. It is the sole donor in which the probiotic genus *Bif*i*dobacterium* engrafted in the mouse colon. In contrast, for the other probiotic-conditioned microbiota recipient mice (donors 13 and 14), both probiotic genera (*Bifidobacterium and Lactobacillus*) were not among the most abundant genera and did not effectively expand in the murine colonic niche.

**Figure 1.**
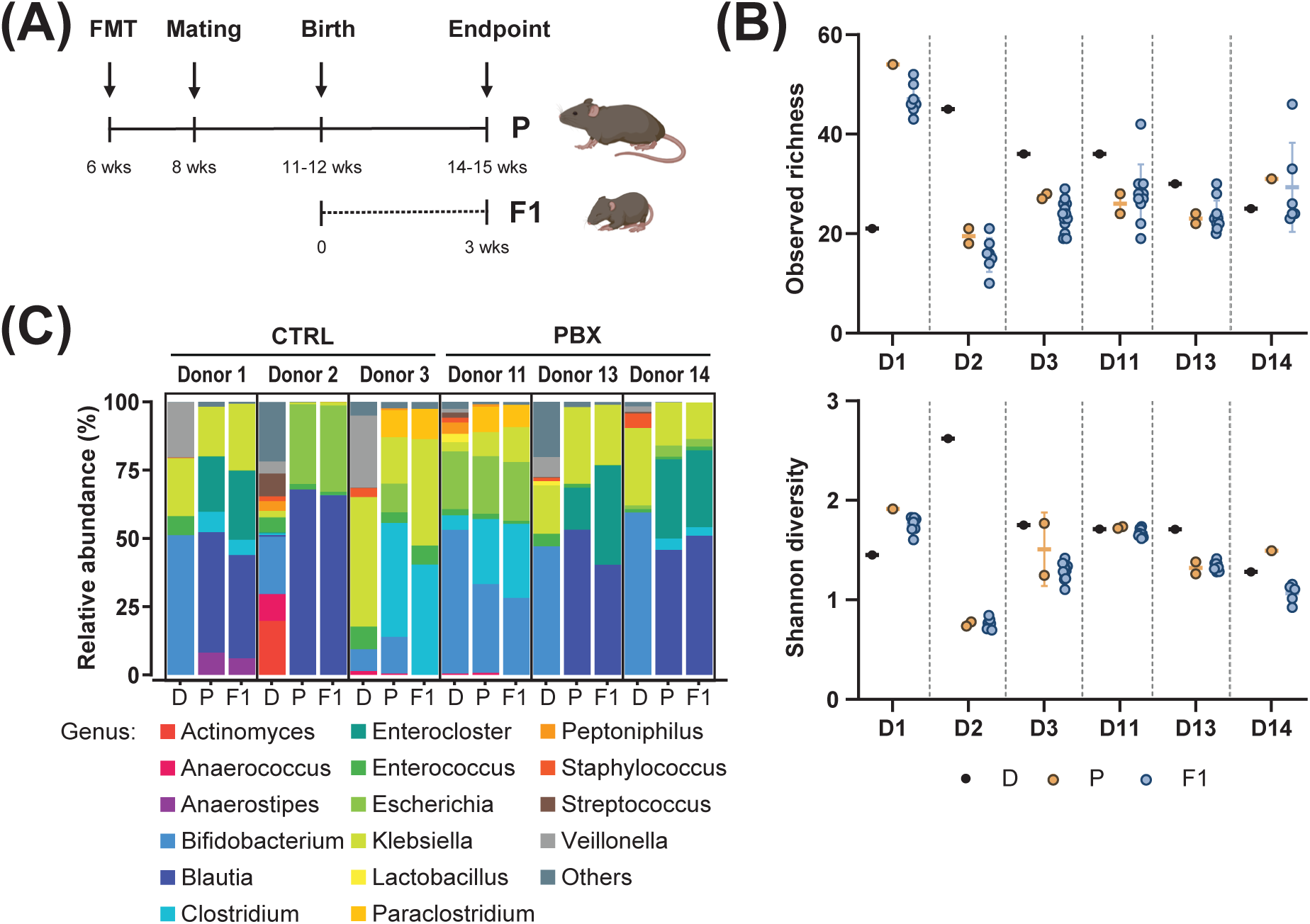
Colonization of mice with fecal microbiota of preterm donors. **(A)** Schematic workflow of experiments. Adult GF mice received fecal microbiota transfer (FMT) from a preterm human donor (D) treated with placebo (CTRL) or probiotics (PBX) via gavage. Animals were mated after 2 weeks. Three weeks after giving birth, adults (P) and pups (F1) were sampled. **(B)** α-diversity as observed amplicon sequence variance (ASV) richness and Shannon diversity. **(C)** Average relative abundance (%) of the most abundant bacterial genera (abundance > 0.5%). Results are pooled from 1 (Donors 1, 11, 13, 14) or 2 (Donors 2, 3) independent experiments.

**Table 1.** Clinical characteristics of the study population.

| Donor | Gestational age (weeks) | Delivery mode | Sample age (days) | Gender | Neonatal probiotic* | Prenatal antibiotic to the mothers** | Postnatal antibiotic to the mothers | Neonatal antibiotic |
| --- | --- | --- | --- | --- | --- | --- | --- | --- |
| D1 | 31+1 | C-Section | 30 | M | No | Yes | No | No |
| D2 | 31+5 | C-Section | 30 | M | No | Yes | No | No |
| D3 | 31+2 | C-Section | 32 | F | No | Yes | No | No |
| D11 | 32+5 | C-Section | 31 | F | Yes | No | No | No |
| D13 | 32+1 | C-Section | 32 | M | Yes | No | No | No |
| D14 | 31+6 | C-Section | 31 | F | Yes | Yes | No | No |

\**Bifidobacterium animalis* subsp. *lactis*, *Bifidobacterium longum* subsp. *infantis*, *Lactobacillus* *acidophilus*
\*\*2 days before birth

### Probiotic-conditioned microbiota alters both adaptive and innate immune responses under homeostasis

A comprehensive analysis of key immune cell populations in 3-week-old pups (F1) born to humanized mothers was conducted using high-dimensional spectral flow cytometry. The analytical approach employed included a panel targeting general and key populations of both innate and adaptive immunity, thereby providing broad coverage of the immune system (**Figure S2**). To obtain a comprehensive perspective on peripheral and local responses, a variety of tissues, including the colon, SI, and spleen, were collected. Most analyzed innate immune cell populations, including neutrophils, macrophages, conventional dendritic cells (cDCs) and innate lymphoid cells (ILCs), were comparable in spleens of mice humanized with feces from probiotic- versus placebo-treated individuals. In contrast, analysis of mucosal organs revealed a significant decrease in most innate immune cell populations in probiotic-humanized mice, with two exceptions: ILCs remained stable in colon and SI, while natural killer (NK) cells increased in SI (**Figure 2A-E**).

**Figure 2.**
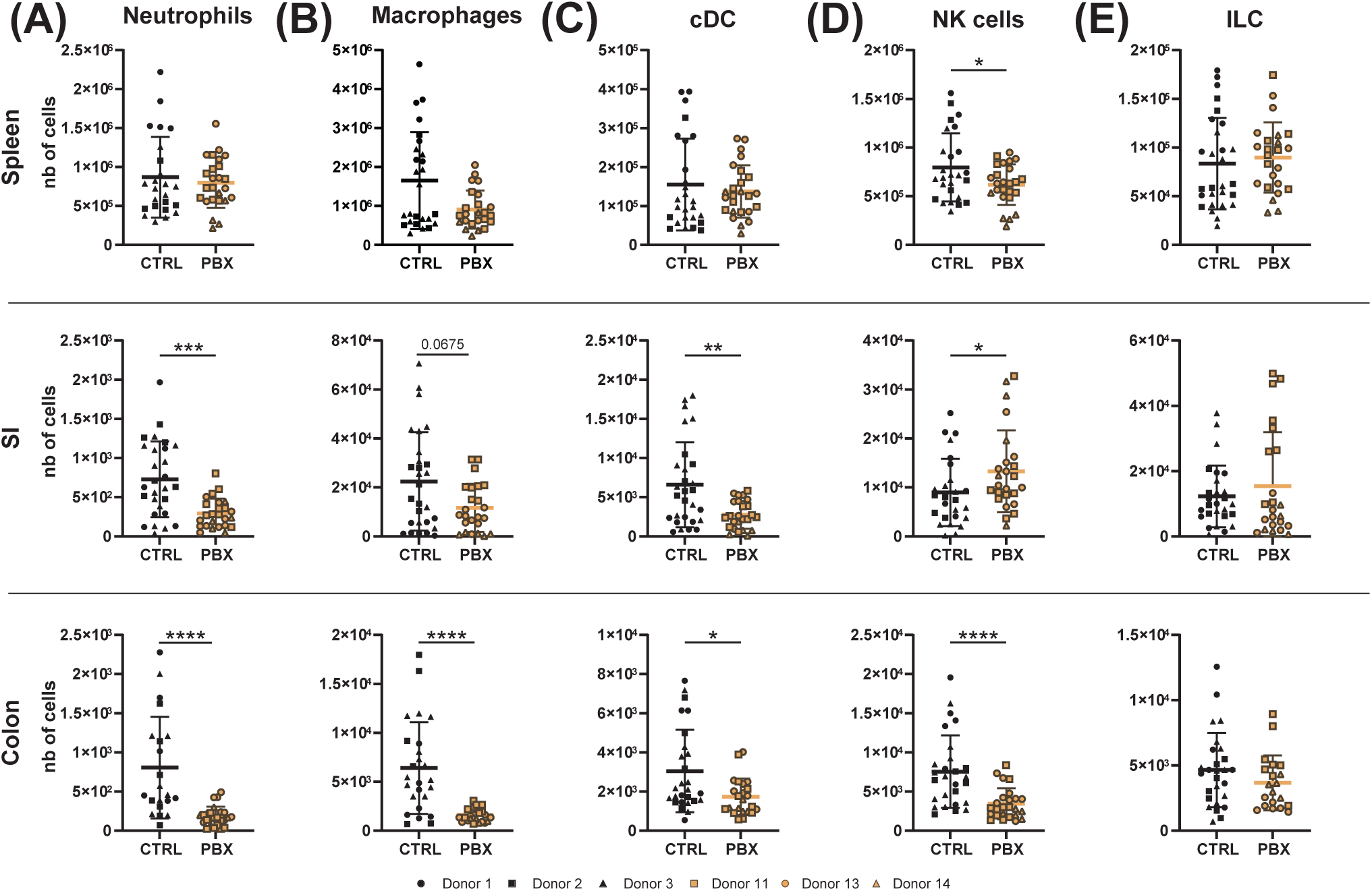
Probiotic-induced microbiota affects innate immune cells in an organ-dependent manner. Spleen, small intestine (SI) and colon of 3-week-old neonates to mothers humanized with placebo (CTRL) or probiotics (PBX) treated preterm feces were collected and analyzed by flow cytometry. **(A)** Cell counts of neutrophils, **(B)** macrophages, **(C)** conventional dendritic cells (cDC), **(D)** NK cells and **(E)** innate lymphoid cells (ILC). Results are pooled from 1 (Donors 1, 11, 13, 14) or 2 (Donors 2, 3) independent experiments and are presented as mean ± SD. Each symbol represents one mouse: CTRL donors - Donor 1 (●, n=8), Donor 2 (▪, n=7), Donor 3 (▴, n=13); PBX donors - Donor 11 (□, n=10), Donor 13 (○, n=10), Donor 14 (△, n=5) (* = p<0.05; ** = p<0.005; **** = p <0.0001).

We hypothesized that the probiotic supplementation would not only affect the innate, but also the adaptive immune system. We observed that the number of conventional CD4^+^ T cells (Tconv) was unchanged in the spleen and SI after humanization with probiotic-treated feces, while it was strongly reduced in the colon, an organ heavily colonized by commensal microorganisms. The same trends were observed for the number of regulatory T cells (Tregs) and CD8^+^ T cells (**Figure 3A**). Regarding cytokine production by Tconv, levels of IFN-γ secretion in spleen and colon increased in probiotic-conditioned animals, while the percentage of IL-17A^+^ Tconv remained constant in all examined organs (**Figure 3B**). Furthermore, we observed an overall decrease in Th1 and Th17 subsets in mice with probiotic-conditioned microbiota, while Th2 cells were significantly increased in the spleen and colon (**Figure S3A**). Additionally, a modest increase in IL-10^+^ Tregs, which represent a small population, was observed in the spleen and colon (**Figure 3C**). An increase in B cells was observed in the systemic immune systems of humanized mice with probiotic-conditioned microbiota, whereas in the colon an opposite trend was observed (**Figure 3D**). Moreover, levels of IgG subclasses (i.e., IgG1, IgG2a and IgG2b) in cecal content were elevated in neonates colonized with the probiotic-conditioned microbiota compared to those with the placebo-conditioned microbiota (**Figure S3B**). In contrast, serum cytokine concentrations remained unchanged, indicating an absence of systemic inflammation (**Figure S3C**).

**Figure 3.**
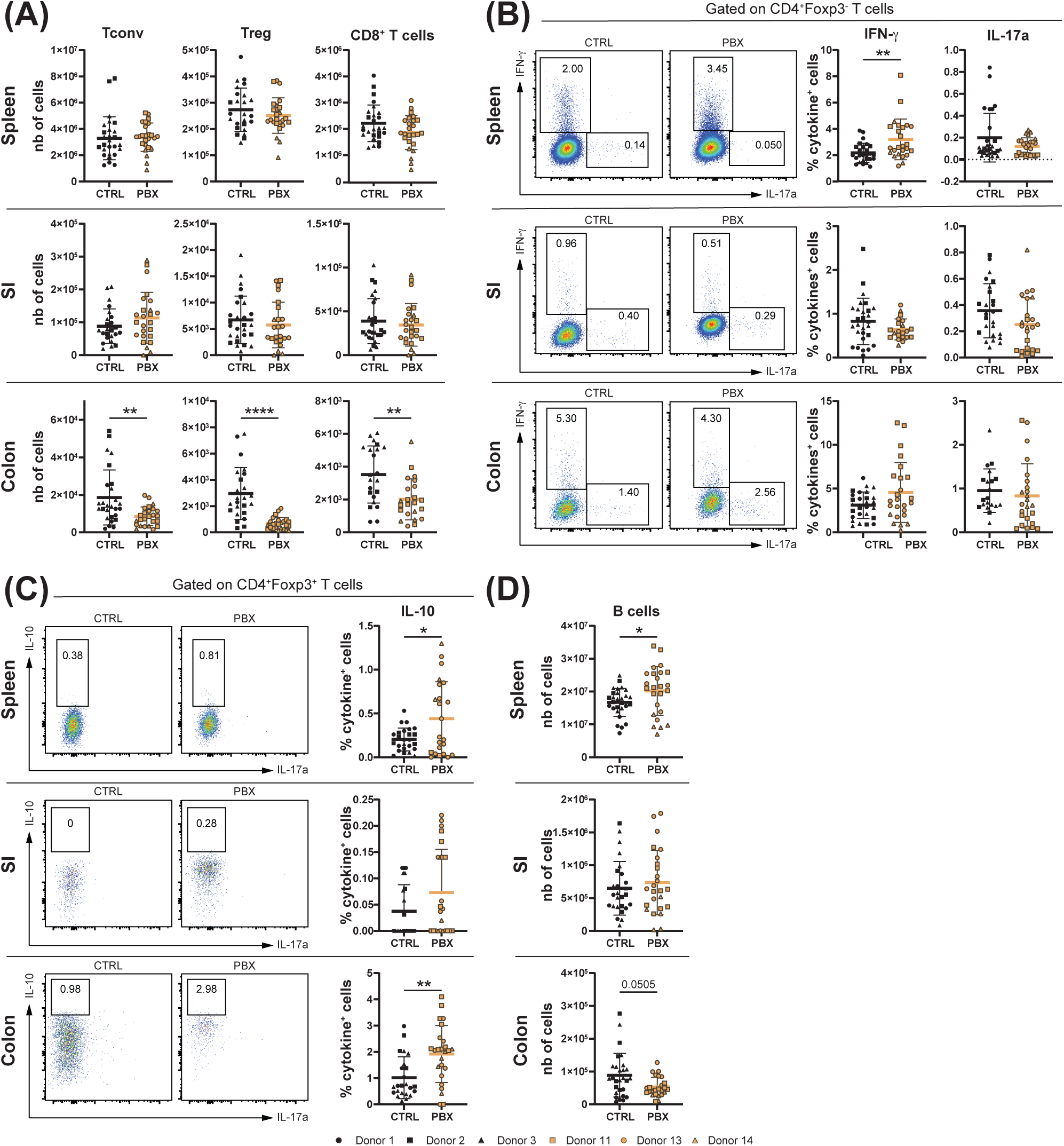
Probiotic-induced microbiota dampens colonic adaptive immunity and promotes IL-10 production. Spleen, small intestine (SI) and colon of 3-week-old neonates born to mothers humanized with placebo (CTRL) or probiotics (PBX) treated preterm feces were collected and analyzed by flow cytometry. **(A)** Cell counts of conventional (Tconv)(CD4^+^Foxp3^-^), regulatory (Treg)(CD4^+^Foxp3^+^) and cytotoxic (CD8^+^) T cells. **(B)** Percentages of Tconv expressing IFN-γ or IL-17A. **(C)** Percentages of Tregs expressing IL-10. **(D)** Cell counts of B cells. Results are pooled from 1 (Donors 1, 11, 13, 14) or 2 (Donors 2, 3) independent experiments and are presented as mean ± SD. Each symbol represents one mouse: CTRL donors - Donor 1 (●, n=8), Donor 2 (▪, n=7), Donor 3 (▴, n=13); PBX donors - Donor 11 (□, n=10), Donor 13 (○, n=10), Donor 14 (△, n=5) (* = p<0.05; ** = p<0.005; **** = p <0.0001).

Our results demonstrate that probiotic-conditioned microbiota comprehensively remodel the immune landscape in humanized mice, affecting both adaptive and innate immune cell populations. Changes include organ-specific modulation of T cell populations with shifts toward anti-inflammatory profiles and increased immunoglobulin production. Concurrently, the majority of innate immune cell populations underwent a decrease in mucosal organs, with the notable exception of stable ILCs and increased NK cells in the SI, highlighting the broad immunomodulatory impact of microbiota modulated by probiotic supplementation.

### Probiotic-induced immune modulation alters susceptibility to gastrointestinal infection

The immunomodulatory effects of the probiotic led us to hypothesize a potential differential impact on susceptibility to gastrointestinal infection. To assess this in a controlled setting, we used EPEC as a well-characterized model pathogen for neonatal enteric infection. Prior work has shown that EPEC colonization in mice depends on age rather than on microbiota composition, and that neonatal infection induces only transient, postnatal dysbiosis (34, 35). Neonates born to mothers humanized with placebo or probiotic-treated preterm feces were infected with 10^7^ CFU of EPEC at postnatal day 5 (**Figure 4A**). Daily monitoring of body weight gain during the first week post-infection served as an indicator of overall health, with a control group receiving the carrier alone (PBS) for comparison. While the probiotic-conditioned microbiota did not affect the development of non-infected mice when compared to the placebo-treated controls, it was detrimental to growth when the mice were infected. Indeed, mice with probiotic-conditioned microbiota gained less body weight over the 7-day period compared to the mice humanized with placebo-treated feces. In the placebo group, we observed that the infection itself had no effect on the bodyweight curve (**Figure 4B**). Seven days post-infection, stomach, liver, SI and colon contents were collected to assess bacterial burden. However, we did not detect differences between the two groups in terms of the number of CFUs present in the organs or contents (**Figure 4C**). Nevertheless, although not significant, a higher percentage of mice from the placebo group eradicated the bacteria after one week, while colonies persisted in a greater proportion of the probiotic-conditioned group (35% versus 32% eradication in the stomach; 42% versus 23% in the SI content; 35% versus 18% in the colon content; 58% versus 54% in the liver) (**Figure 4D**). These findings suggest that while probiotic treatment modulates immune responses, it may potentially influence susceptibility to gastrointestinal infections under certain conditions.

**Figure 4.**
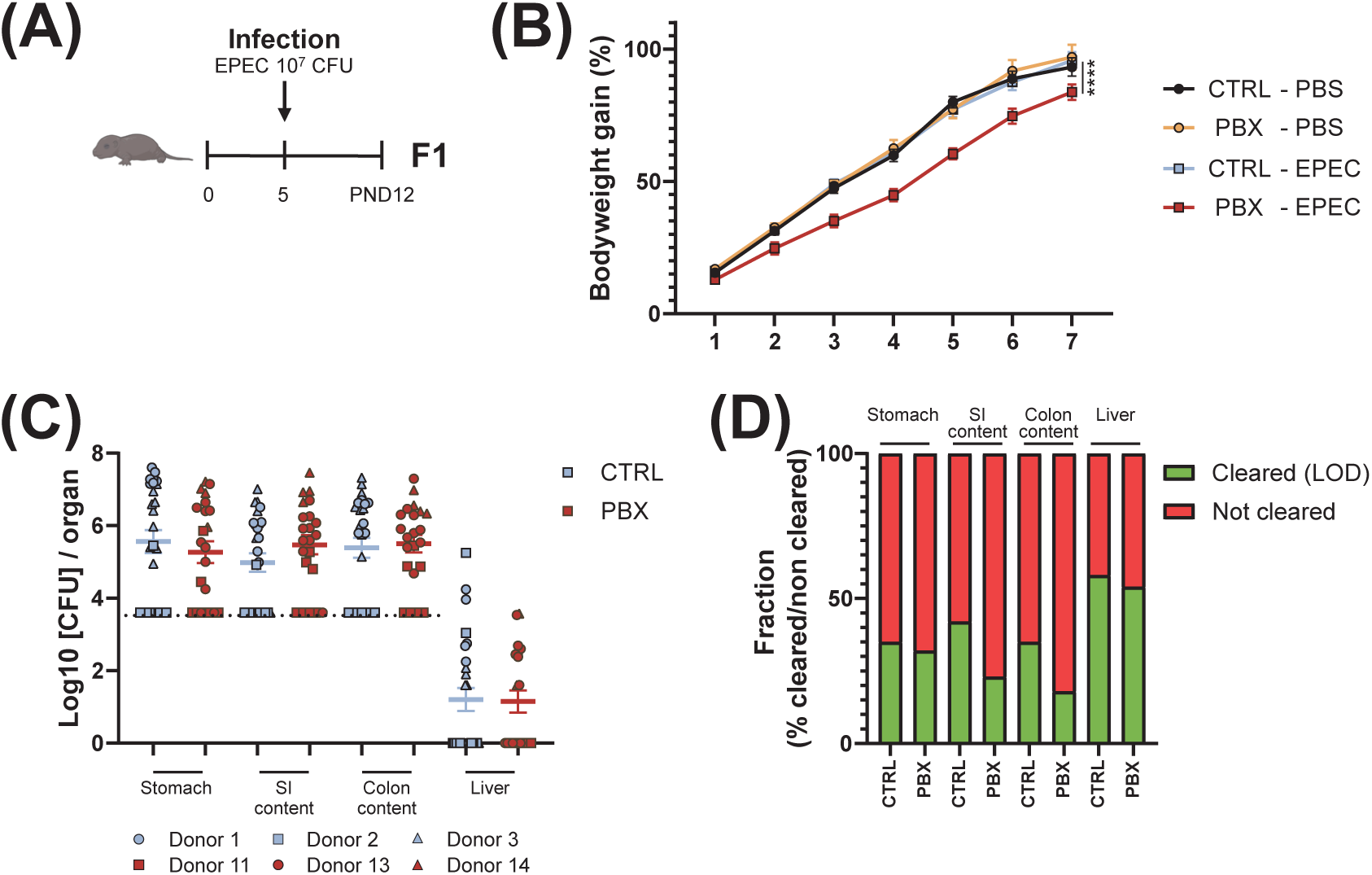
Probiotic-conditioned microbiota influences susceptibility to infection. **(A)** Schematic workflow of experiments. Adult GF mice received fecal microbiota transfer (FMT) from placebo (CTRL) or probiotic (PBX) treated preterm human donor via gavage. Animals were mated after 2 weeks. Five days after birth, the pups (F1) were infected via intragastric injection with 10^7^ CFU of EPEC or with the carrier alone (PBS) and were sampled 7 days post-infection. **(B)** Gain in body weight calculated as a percentage of pre-infection weight. **(C)** The indicated tissues and feces were collected, homogenized, and plated. Colony-forming units (CFU) were counted and expressed as CFU per organ or content. The dashed line represents the limit of detection (LOD). **(D)** Percentages of mice that cleared the bacteria (determined by the LOD) in each organ or content. Results are pooled from 1 (Donors 1, 11, 14), 2 (Donors 2, 13) or 3 (Donor 3) independent experiments and are presented as mean ± SEM. Each symbol represents one mouse. Sample sizes per donor are given as *n*PBS/*n*EPEC: CTRL donors - Donor 1 (●, 3/7), Donor 2 (▪, 8/9), Donor 3 (▴, 7/10); PBX donors - Donor 11 (□, 7/5), Donor 13 (○, 11/11), Donor 14 (△, 0/6) (**** = p <0.0001).

### Downregulation of the immune response to EPEC in probiotic-conditioned mice may contribute to delayed resolution of infection

Since the innate immune cell compartment was strongly altered at 3 weeks of age under homeostasis, we sought to determine if this was also the case at 12 days of age, both under homeostasis and after subjecting the offspring to EPEC challenge. Under homeostatic conditions, only the number of cDCs in the SI was significantly reduced after mice were colonized with probiotic-treated feces when compared to placebo-treated controls, and for NK cells a trend towards fewer cells was observed in the same organ (**Figure 5C**). All other innate immune cell subsets showed comparable numbers between the two groups. Upon infection, however, the number of neutrophils (**Figure 5A**), macrophages (**Figure 5B**), cDCs (**Figure 5C**), NK cells (**Figure 5D**), and ILCs (**Figure 5E**) decreased significantly in the spleens of the probiotic group. In the SI, only ILC numbers decreased significantly in the probiotic group compared to the placebo group. For all other innate immune cell populations, only a trend toward reduced numbers was observed (**Figure 5E**).

**Figure 5.**
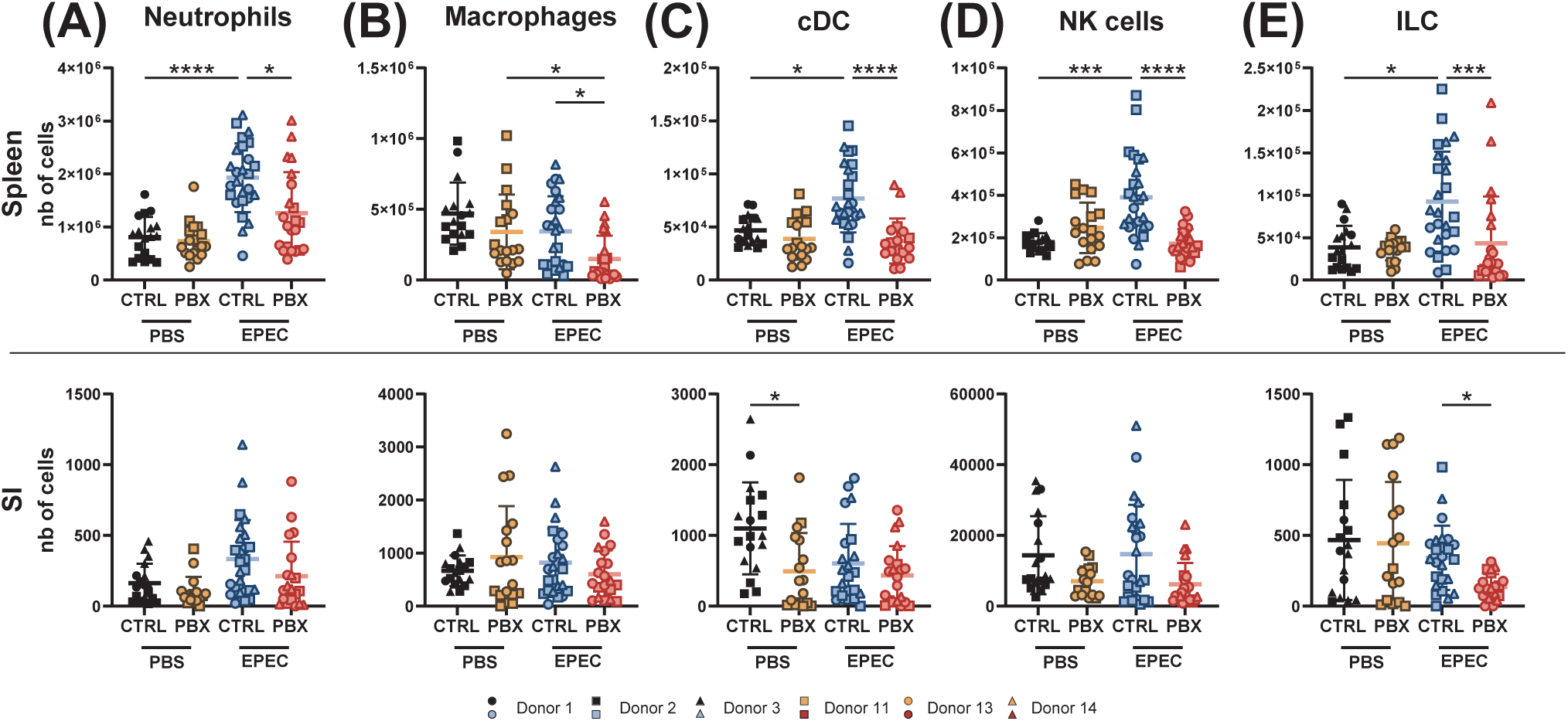
Probiotic-induced microbiota negatively affects the systemic innate response during early life gastrointestinal infection. Spleens and small intestines (SI) of sham (PBS) or infected (EPEC) neonates at postnatal day 12, born to mothers humanized with placebo (CTRL) or probiotics (PBX), were collected 7 days post-infection and stained for FACS analysis. **(A)** Cell counts of neutrophils, **(B)** macrophages, **(C)** conventional dendritic cells (cDC), **(D)** NK cells, and **(E)** innate lymphoid cells (ILC). Results are pooled from 1 (Donors 1, 11, 14), 2 (Donors 2, 13) or 3 (Donor 3) independent experiments and are presented as mean ± SD. Each symbol represents one mouse. Sample sizes per donor are given as *n*PBS/*n*EPEC: CTRL donors - Donor 1 (●, 3/7), Donor 2 (▪, 8/9), Donor 3 (▴, 7/10); PBX donors - Donor 11 (□, 7/5), Donor 13 (○, 11/11), Donor 14 (△, 0/6) (* = p<0.05; *** = p<0.001; **** = p <0.0001).

We next examined the adaptive response to EPEC infection in systemic (spleen) and mucosal (SI) organs 7 days post-infection, at postnatal day 12. The adaptive compartment in the SI was not affected by either infection or probiotic-conditioned microbiota, similar to what was observed at 3 weeks of age. However, upon infection, adaptive immunity was impaired in spleens of mice humanized with probiotic-treated feces. In these mice, we observed decreased numbers of Tconv, Tregs, and CD8^+^ T cells (**Figure 6A**), and also B cells (**Figure S4A**). However, the distribution of Th1 and Th17 subsets was unchanged when comparing the placebo and probiotic groups (**Figure 6B**). Interestingly, although we did not observe a change in the absolute number of T cells in the SI between uninfected and infected groups, we demonstrated that mice could mount a T cell response even at this early time point. This was evidenced by an increase in the proportion of Th1 and Th17 cells in both spleen and SI upon infection (**Figure 6B**) and an increase in the percentage of IL-17A-producing Tconv in the spleen and also IFN-γ and IL-17A-producing Tconv in the SI (**Figure 6C**). The increase in IL-17A^+^ cells upon infection was also observed for Tregs in both the spleen and SI, while a significant increase in the frequency of IL-10^+^ Tregs upon infection was only found in the SI of mice from the probiotic group (**Figure 6D**). Increased IL-10 production upon infection was also observed in B cells from the probiotic group (**Figure S4B**). However, we did not observe any significant changes in the secretion of immunoglobulins in the cecal contents upon infection (**Figure S4C**).

**Figure 6.**
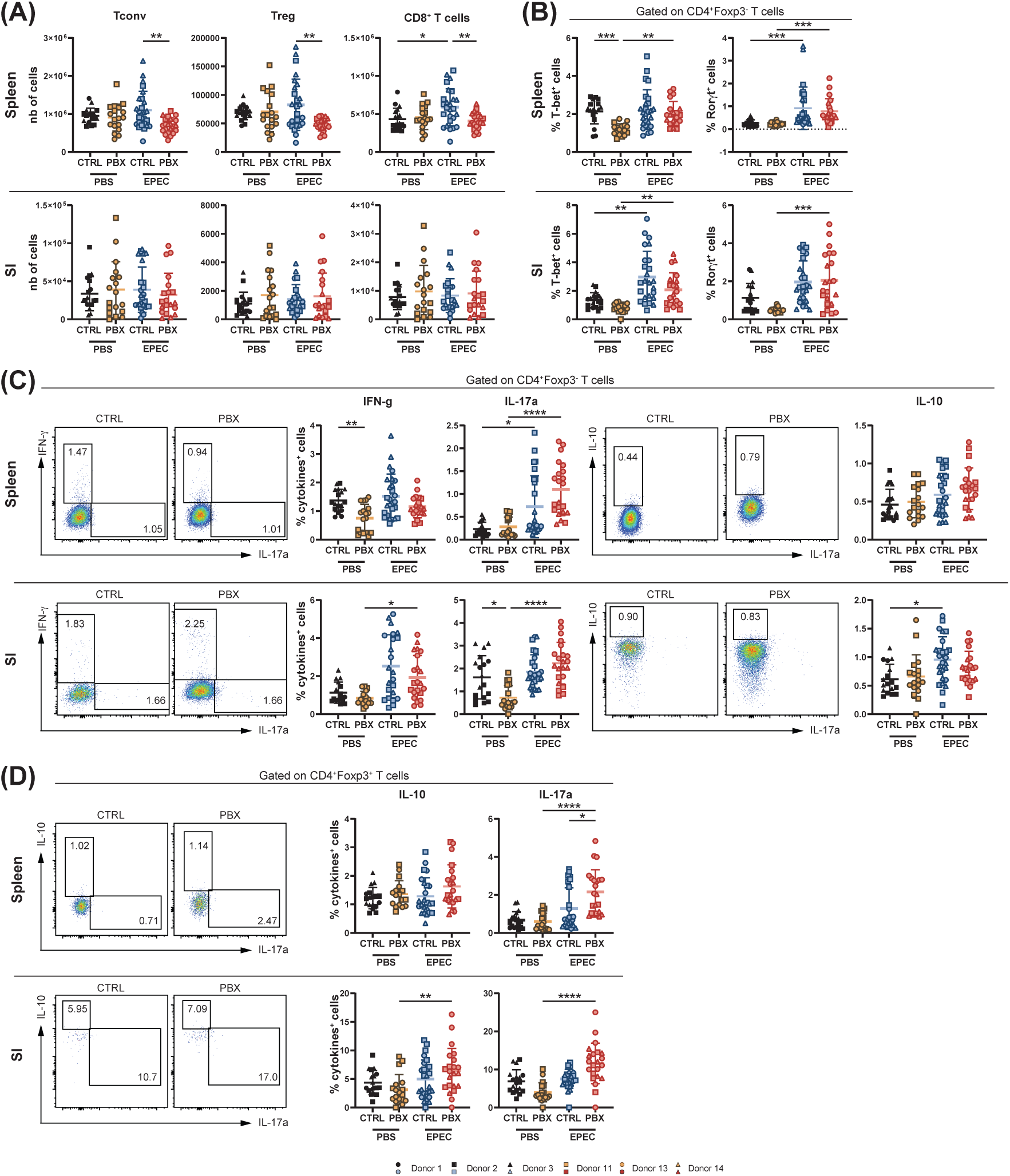
Probiotic-conditioned microbiota downregulate systemic adaptive immunity under early life gastrointestinal infection. Spleen and small intestine (SI) of sham (PBS) or infected (EPEC) neonates at postnatal day 12 born to mothers humanized with placebo (CTRL) or probiotics (PBX) treated preterm feces were collected 7 days post-infection and analyzed by flow cytometry. **(A)** Cell counts of conventional (Tconv)(CD4^+^Foxp3^-^), regulatory (Treg)(CD4^+^Foxp3^+^) and cytotoxic (CD8^+^) T cells. **(B)** Percentage of Tconv expressing the transcription factors associated with Th1 (T-bet) or Th17 (Rorγt). **(C)** Percentages of Tconv expressing IFN-γ, IL-17A or IL-10. **(D)** Percentages of Tregs expressing IL-10 or IL-17A. Results are pooled from 1 (Donors 1, 11, 14), 2 (Donors 2, 13) or 3 (Donor 3) independent experiments and are presented as mean ± SD. Each symbol represents one mouse. Sample sizes per donor are given as *n*PBS/*n*EPEC: CTRL donors - Donor 1 (●, 3/7), Donor 2 (▪, 8/9), Donor 3 (▴, 7/10); PBX donors - Donor 11 (□, 7/5), Donor 13 (○, 11/11), Donor 14 (△, 0/6) (* = p<0.05; ** = p<0.005; *** = p <0.001; **** = p <0.0001).

In conclusion, probiotic-conditioned microbiota impaired splenic adaptive immunity during EPEC infection while maintaining functional T cell responses. Most critically, it caused decreased innate immune cell populations in the gut and systemically in the spleen, potentially explaining the lower bacterial clearance and reduced weight gain observed in mice humanized with probiotic-conditioned microbiota. These findings suggest that while probiotics influence immune development, they may also alter resistance to infections in early life.

## Discussion

Understanding how early-life microbial environments shaped by probiotic supplementation influence the age-dependent programming of innate and adaptive immunity, particularly in the highly vulnerable preterm population, remains incomplete. To address this gap in the current study and in future studies, we developed a murine neonatal model system that replicates key aspects of the preterm microbiota, albeit with limitations in microbial richness and diversity. Our findings revealed that microbiota conditioned by early-life probiotic supplementation distinctly shaped both the neonatal adaptive and innate immune systems in 3-week-old mice. We observed marked, organ-dependent reductions in adaptive immune cells, with the colon being more affected than the SI and spleen, while innate immune compartments in both the SI and colon also exhibited significant alterations. These microbiota-mediated effects were both organ- and age-dependent, as no significant immune differences were detected under homeostatic conditions at postnatal day 12 in the SI or spleen. Furthermore, during gastrointestinal infection, mice receiving probiotic-conditioned microbiota recovered more slowly, as reflected by delayed weight gain and reduced bacterial clearance. is increased susceptibility may result from impaired coordination of innate and adaptive responses, potentially facilitating pathogen persistence and dissemination. Collectively, these findings provide insight into potential mechanisms by which postnatal probiotic supplementation might influence infection outcomes in preterm infants. They underscore the complex interplay between microbial colonization and host defense mechanisms, highlighting the intricate nature of early-life immune system development and the need for careful consideration of probiotic interventions in vulnerable neonatal populations.

Despite its limitations, using GF mice as recipients for FMT offers the advantage of working with genetically identical organisms, where only the gut microbiota composition differs. Moreover, we had the unique opportunity to access preterm fecal samples from the PRIMAL trial, a RCT that studied the impact of probiotic intervention on limiting colonization with pathobionts and shaping the gut microbiome in early life (25). For this study, we used three fecal samples from placebo-treated preterm infants and three fecal samples from probiotic-treated preterm infants to perform the FMT. While this sample size limits generalizability and precludes assessment of donor-specific effects, the consistency of our findings across donors suggests robust biological effects of probiotic-conditioned microbiota. Future studies with larger donor cohorts are needed to validate these observations and explore potential inter-individual variability. Although we did not observe effective engraftment of probiotic species in mice, we could clearly demonstrate effects with the probiotic-conditioned microbiota that were not donor-dependent. The heterogeneity observed in our results was not attributed to donor dependency but rather to biological variability, environmental factors, developmental differences, or technical variability inherent to the experimental system. Studies utilizing humanized models often do not report engraftment efficiency compared to their original human samples. In the present study, we observed a transfer rate of approximately 64% (ranging from 55 to 78%, depending on the donor) of OTUs at the genus level from preterm infants to the murine parental generation and noted variations in the expansion of specific bacterial genera within the gut environment of our established model. Other studies have shown that approximately 85–88% of human gut microbial genera can successfully colonize GF mice within seven days post-FMT (36–39). However, this engraftment rate decreases over time, with about 52–58% of the donor microbiota persisting after one week following the last FMT dose and maintaining stability for the subsequent three weeks (38). The timing, colonization duration, and dosage are crucial parameters for establishing gnotobiotic mice (37, 40). To maximize mimicry of physiological birth when colonization begins, we colonized the parent population rather than the F1 generation, ensuring microbial exposure directly at birth. Early-life exposure is crucial for establishing intestinal homeostasis and mitigating the immunological defects of GF mice, which, for some parameters, can only be restored within this critical window (40). As previously reported by Lundberg et al. (39), we observed vertical transmission of the gut microbiota from mother to F1 generation. They demonstrated that anti-inflammatory bacteria from humans, such as *Bifidobacterium*, do not colonize the mouse gut in a C57BL/6 background. When comparing the relative abundance in humanized mice to their original donors, similar differences were observed in both parental and F1 animals, with the genera *Blautia* and *Lachnospiraceae* dominating the mouse gut (39). These findings align closely with our observations, where *Blautia* and *Enterocloster* were the most prevalent genera in the mouse colon, followed by *Klebsiella*. However, we also detected engraftment of *Bifidobacterium* in the recipient mice from donor 11 across generations and in the parental generation from donor 3. These two exceptions highlight the variability in microbiota engraftment and demonstrate that donor-specific factors can override general colonization patterns. This suggests that certain donor microbiota compositions or specific *Bifidobacterium* strains may be more adaptable to the mouse gut environment, or that particular donor-recipient combinations create more favorable conditions for engraftment of typically non-colonizing species. These intrinsic differences between human and mouse intestines, including immune system maturity and gut environment, can influence colonization efficiency, with certain human gut microbes undergoing significant changes in relative abundance after transplantation into GF mice and only 47% of human gut microbes being re-established at the species level (41). While donor samples clustered more tightly together than their corresponding mouse recipients, indicating incomplete recapitulation of the donor microbial community structure, the transferred microbiota retained key functional characteristics that distinguished probiotic-conditioned from placebo-conditioned groups. This suggests that despite structural differences in community composition, functionally relevant microbial signatures were successfully transferred and maintained in the humanized mice. Importantly, the increased richness and diversity observed in the gnotobiotic mice, as well as the detection of taxa absent from donor sequencing profiles, can also reflect the outgrowth of low-abundance bacteria that were present in the original samples but below the detection threshold. Such expansion of rare taxa is well documented in human-to-mouse FMT models, where the low-competition environment of GF mice allows minor community members to proliferate (42). This possibility should therefore be considered alongside technical factors when interpreting differences between donor and recipient microbial profiles. Moreover, as mentioned earlier, the duration of colonization is an important parameter that influences engraftment efficiency. Here, we colonized the dams by a single gavage, 2 weeks before mating. However, it has been demonstrated that multiple gavages significantly increase the number of detected taxa and lead to a composition more similar to the donor microbiota while reducing inter-animal variance (43).

Both the adaptive and innate immune system were impacted by the probiotic-conditioned microbiota in 3-week-old mice, but in an organ-dependent manner. As described in the PRIMAL RCT, preterm infants given probiotics tend to develop a less inflammatory microbiota (24). Consequently, we demonstrated that when this altered microbiota is transferred to mice, it reduces immune activation, with fewer immune cells accumulating in the colon, as evidenced by the significantly reduced numbers of neutrophils, macrophages, cDCs, and NK cells. These cells, with the exception of NK cells, were also reduced in the SI, indicating decreased recruitment of cells and therefore favoring a tolerogenic environment. Furthermore, all major adaptive immune cell populations were decreased in the colon, while the spleen and SI remained unaffected. By definition, the spleen is not colonized by the gut microbiota, and microbiota-dependent activation of splenic cells can only occur through indirect stimulation via metabolites or bacterial antigens (9). In contrast, both the colon and the small intestine are colonized organs. However, the colon is the primary site where microbial communities from an FMT tend to establish, whereas the SI is more hostile to colonization due to faster transit, bile acids, and oxygen exposure. Therefore, microbial changes and subsequent immune effects are more pronounced in the colon of recipient mice (44). We demonstrated that the numbers of B cells, CD8^+^ and CD4^+^ T cells, comprising both Tconv and Tregs, were all reduced in the colon of probiotic-treated mice when compared to placebo-treated controls. The probiotic strains used in the PRIMAL RCT are known to enhance mucosal barrier function, stimulate Tregs, and promote immune tolerance, especially in the colon (45, 46). Indeed, we showed an increased number of IL-10^+^ Tregs. Interestingly, this observation was made in both the colon and the spleen. In addition to the expansion of IL-10-producing cells, we also observed a decreased proportion of Th1 and Th17 cells. Thus, introducing the probiotic-conditioned microbiota in GF mice rebalances the immune environment in the colon toward a tolerogenic, non-inflammatory state, resulting in a reduced need for immune surveillance across both adaptive and innate immune cells. Collectively, these findings demonstrate that probiotic-conditioned microbiota establishes a systemic shift toward immune tolerance, with the most pronounced effects occurring in directly colonized mucosal sites. Indeed, the localized increase in IgG1 and IgG2 subclasses in the cecum occurred in the absence of systemic cytokine elevation. This suggests a mucosal immune adaptation potentially driven by microbiota shifts, without concurrent systemic immune activation. The IgG1/IgG2 increase supports the hypothesis of a shift toward anti-inflammatory Th2 responses relative to Th1 responses in mucosal tissues (47).

This overall mucosal tolerance may explain the dampened growth of the pups when exposed to pathogenic challenge 5 days after birth. Indeed, mice from the probiotic-conditioned microbiota group showed reduced body weight gain in the week following infection. Although the organ burden of EPEC bacteria was similar between the two groups, we observed lower clearance in the probiotic-conditioned microbiota mice compared to the placebo-conditioned microbiota mice. The observed differences in growth patterns and the trends toward lower bacterial clearance rates in the probiotic-conditioned group compared to the placebo group could indicate a complex relationship between probiotic-induced immune modulation and host response to enteric infections that warrants further investigation. As discussed above, the tolerogenic microbiota environment biases the host immune system toward anti-inflammatory Th2 responses, albeit at the cost of reduced Th1/Th17-mediated bacterial clearance, resulting in impaired host defense against pathogens such as EPEC. To understand how this altered immune landscape responds to sustained pathogenic challenge, we analyzed immune cell populations and functions 7 days post-infection.

In contrast to observations in 3-week-old mice, we did not observe any impact of probiotic-conditioned microbiota on adaptive or innate immune cell populations under homeostatic conditions at postnatal day 12, suggesting an age-dependent effect. The only affected cells at this age were intestinal cDCs. Due to experimental sampling limitations, we were unable to collect sufficient colonic cells at this age to provide accurate flow cytometric analysis and comparison between early and late time points. The innate immune system serves as the first line of defense against enteric bacterial infections. During EPEC infection, multiple innate immune cell types coordinate the initial response: neutrophils act as first responders and migrate locally to the infection site; macrophages engulf and eliminate bacteria through phagocytosis; DCs bridge innate and adaptive immunity by sampling EPEC antigens and presenting them to T cells for activation; NK cells enhance macrophage and DC functions; and ILC reinforce the gut barrier and enhance antimicrobial defense (48–51). Notably, all these innate immune cell populations were reduced in the spleens of infected mice with probiotic-conditioned microbiota, whereas in the SI, only ILC numbers were significantly reduced. This differential pattern between systemic and mucosal compartments may reflect altered immune cell trafficking or recruitment rather than simply local immune suppression, potentially due to probiotic-induced changes in chemokine gradients or immune cell priming during the critical developmental window. Such alterations in innate immune cell availability could contribute to prolonged bacterial persistence and increased mucosal damage, potentially explaining the delayed weight gain observed in mice with probiotic-conditioned microbiota.

Although no differences in the adaptive cell compartment were evident in the non-infected groups at postnatal day 12, infection reduced T and B cell numbers in the spleen of mice with probiotic-conditioned microbiota. Nevertheless, these mice mounted comparable adaptive responses to controls, including increased Th1 and Th17 cells and enhanced IFN-γ and IL-17A production. A closer examination of the Treg subsets revealed an increased number of IL-17A-expressing cells in mice receiving probiotic-conditioned microbiota. These Th17-like Tregs can contribute to host defense; however, an increased proportion may indicate poor regulation of bacterial infections, tissue damage, and loss of immune balance (52). When present in excessive proportions, this indicates regulatory stress and suggests that the system is shifting toward a Th17-dominant state (53). A recent study demonstrated that the specific deletion of the IL-23 receptor in Tregs decreases their conversion into Th17-like phenotypes and enhances resistance to *Citrobacter rodentium* infection, indicating that excessive Th17-like Tregs correlate with dysregulated immunity during enteric infections (54). Although not essential for survival during EPEC infection, B cells play a role in bacterial clearance (55). Here, we observed that EPEC infection led to a reduction in B cell numbers in both the SI and spleen of probiotic-conditioned microbiota mice, accompanied by an increased proportion of regulatory B cells producing IL-10. Notably, cecal immunoglobulin levels remained unchanged despite these cellular alterations. This shift in B cell composition may impair effective bacterial clearance despite preserved antibody levels, potentially contributing to diminished infection control.

Overall, these findings highlight a potential trade-off where probiotic-conditioned immune tolerance, while beneficial under homeostatic conditions, may become detrimental during pathogenic encounters. Whether this represents impaired immune cell development, defective recruitment mechanisms, or functional suppression of existing cells warrants further investigation.

## Abbreviations

cDC – conventional dendritic cell; CFU – colony-forming units; EPEC – Enteropathogenic *Escherichia coli*; FMT – fecal microbiota transplantation; GF – germ-free; IL – interleukin; ILC – innate lymphoid cell; LOD – limit of detection; NK – natural killer; OUT – operational taxonomic units; PRIMAL - priming immunity at the beginning of life; RCT – randomized control trial; SI – small intestine; Tconv – conventional CD4^+^ T cells; Th – T helper cell; Treg – regulatory T cell

## Author Contributions

JS: Conceptualization, Formal analysis, Investigation, Methodology, Visualization, Writing – original draft; TRL: Data curation, Formal analysis; LH: Conceptualization, Investigation, Writing – review & editing; DO: Methodology; MZ: Methodology; FK: Investigation, Writing – review & editing; SP: Data curation, Resources; MW: Data curation, Resources; CH: Data curation, Resources; CF: Formal analysis; NT: Conceptualization; DV: Conceptualization, Data curation, Funding acquisition, Resources, Supervision, Writing – review & editing; TS: Conceptualization, Data curation, Funding acquisition, Supervision, Writing – review & editing; JH: Conceptualization, Funding acquisition, Project administration, Supervision, Writing – review & editing

## Acknowledgments

We acknowledge the Flow Cytometry and Cell Sorting platform, the Genome Analytics facility and the Animal facility at the Helmholtz Centre for Infection Research. We thank Kerstin Beushausen, Jana Keil, Achim Gronow, Martina Palatella and Svetlina Khayat for their technical support.

## Data Availability Statement

The raw 16S rRNA amplicon sequence reads reported in this study have been deposited in the ENA (European Nucleotide Archive) database under accession code PRJEB96747 (https://www.ebi.ac.uk/ena/browser/view/PRJEB96747).

## Supplementary Figures

**Figure S1.**
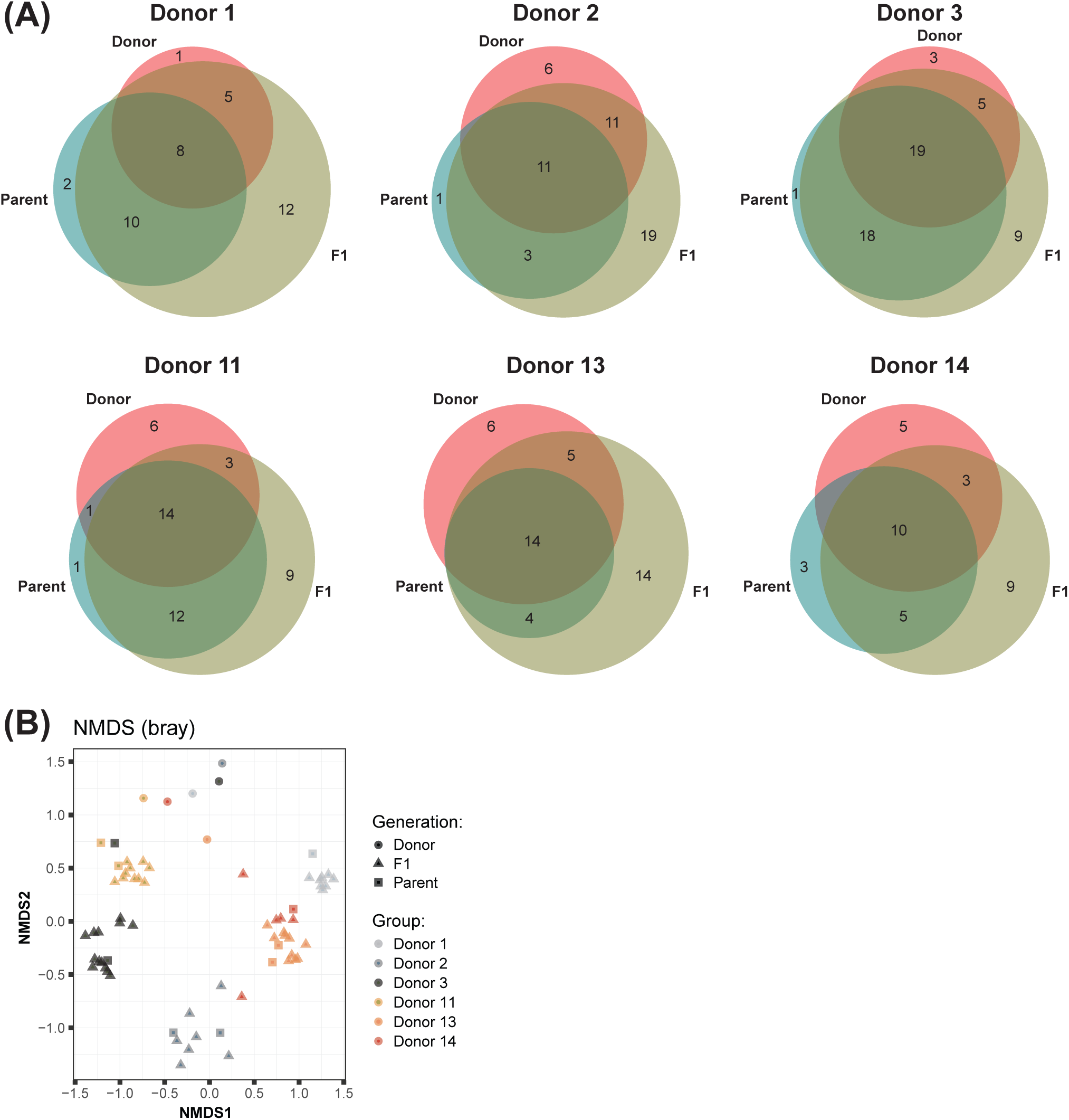
Colonization of mice with different preterm microbiota. **(A)** Venn diagrams displaying shared species between the donor (red), the parent population (teal) and the F1 generation (olive). **(B)** β-diversity of fecal samples from donor, parent and F1 generation was analyzed using the Bray-Curtis dissimilarity matrix and NMDS. Results are pooled from 1 (Donors 1, 11, 13, 14) or 2 (Donors 2, 3) independent experiments.

**Figure S2.**
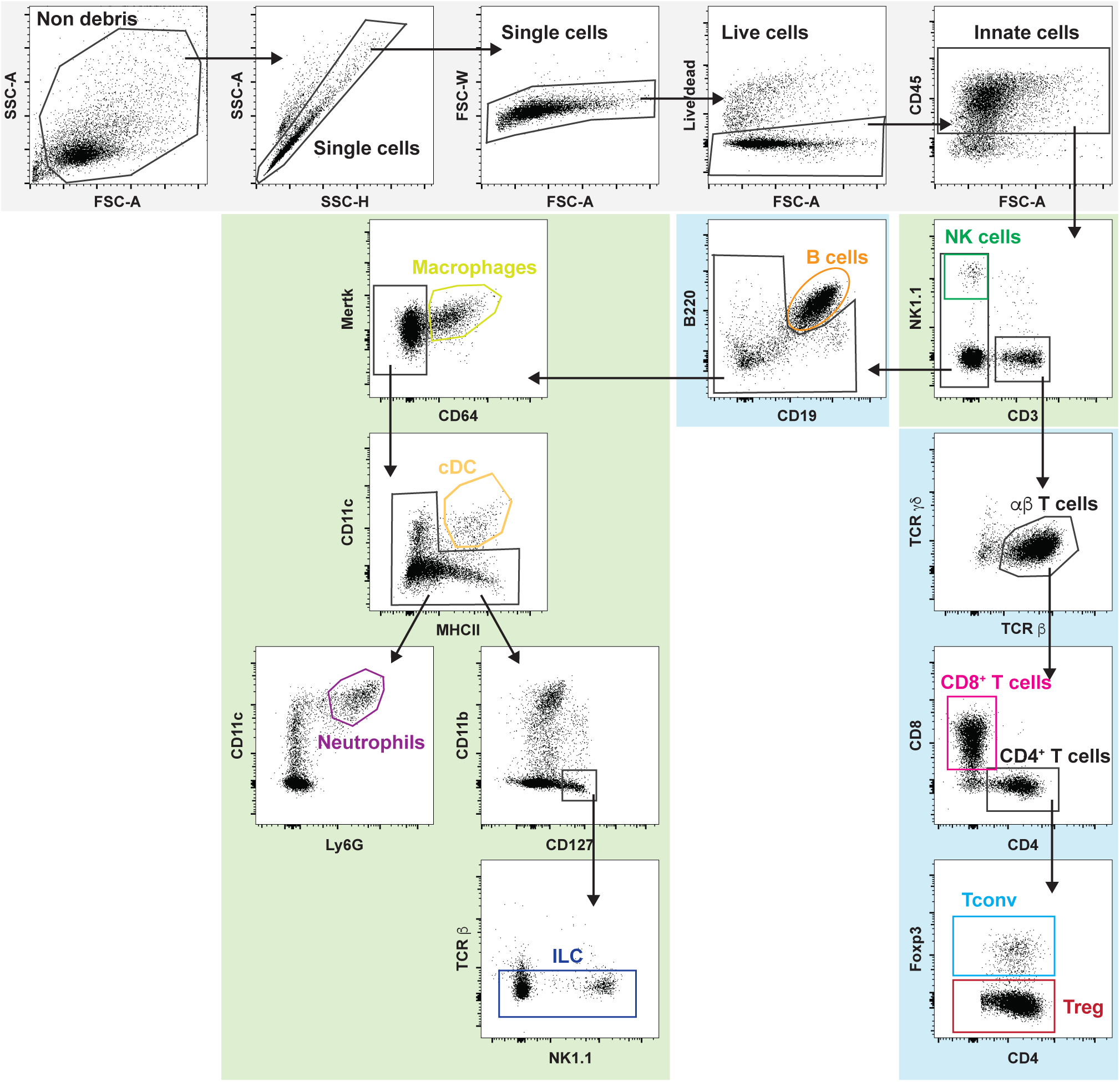
Gating strategy. Representative dot plots to identify the major innate (green box; natural killer (NK) cells, macrophages, conventional dendritic cells (cDC), neutrophils, and innate lymphoid cells (ILC)) and adaptive (blue box, B cells, CD8^+^ T cells, regulatory T cells (Treg), and conventional T cells (Tconv)) immune cells, shown here for 3-week-old spleen.

**Figure S3.**
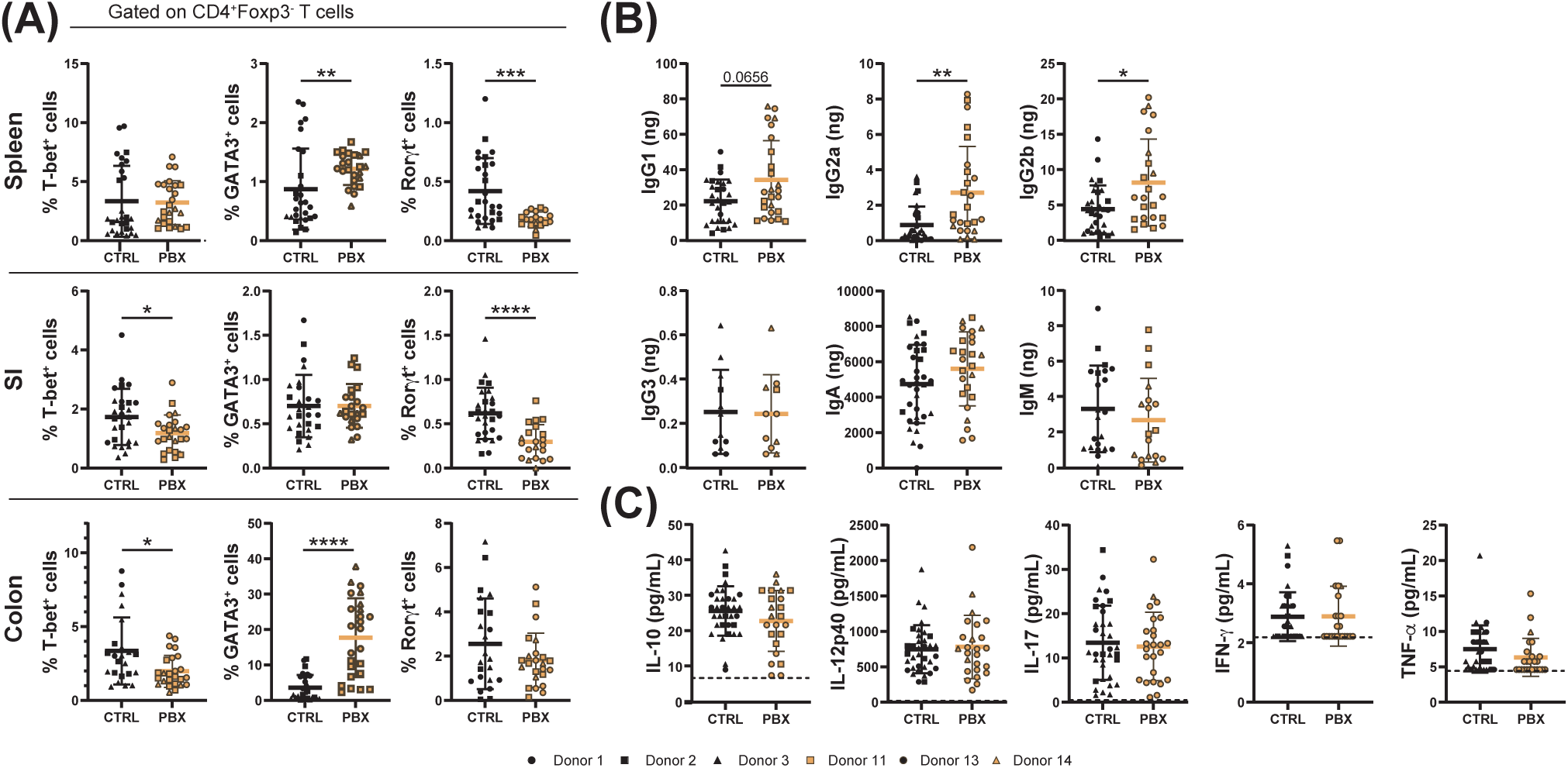
Probiotic-induced microbiota skews T helper differentiation and triggers a selective mucosal IgG subclasses elevation in the absence of systemic inflammation. Spleen, small intestine (SI), colon, cecal content and serum were collected from 3-week-old neonates born to mothers humanized with either placebo (CTRL) or probiotic (PBX)-treated preterm fecal microbiota. The organs samples were analyzed by flow cytometry while the cecal contents and serum were analyzed by multiplex assay. **(A)** Percentages of CD4^+^ conventional T cells (Tconv) expressing the transcription factors associated with Th1 (T-bet) or Th17 (Rorγt). **(B)** Amount of IgG1, IgG2a, IgG2b, IgG3, IgA, and IgM in the total cecal content. **(C)** Concentration of serum cytokines IL-10, IL-12p40, IL-17A, IFN-γ, and TNF-α. The dashed line represents the limit of detection (LOD). Results are pooled from 1 (Donors 1, 11, 13, 14) or 2 (Donors 2, 3) independent experiments and are presented as mean ± SD. Each symbol represents one mouse: CTRL donors - Donor 1 (●, n=8), Donor 2 (▪, n=7), Donor 3 (▲, n=13); PBX donors - Donor 11 (□, n=10), Donor 13 (○, n=10), Donor 14 (△, n=5) (* = p<0.05; ** = p<0.005; *** = p<0.001; **** = p <0.0001).

**Figure S4.**
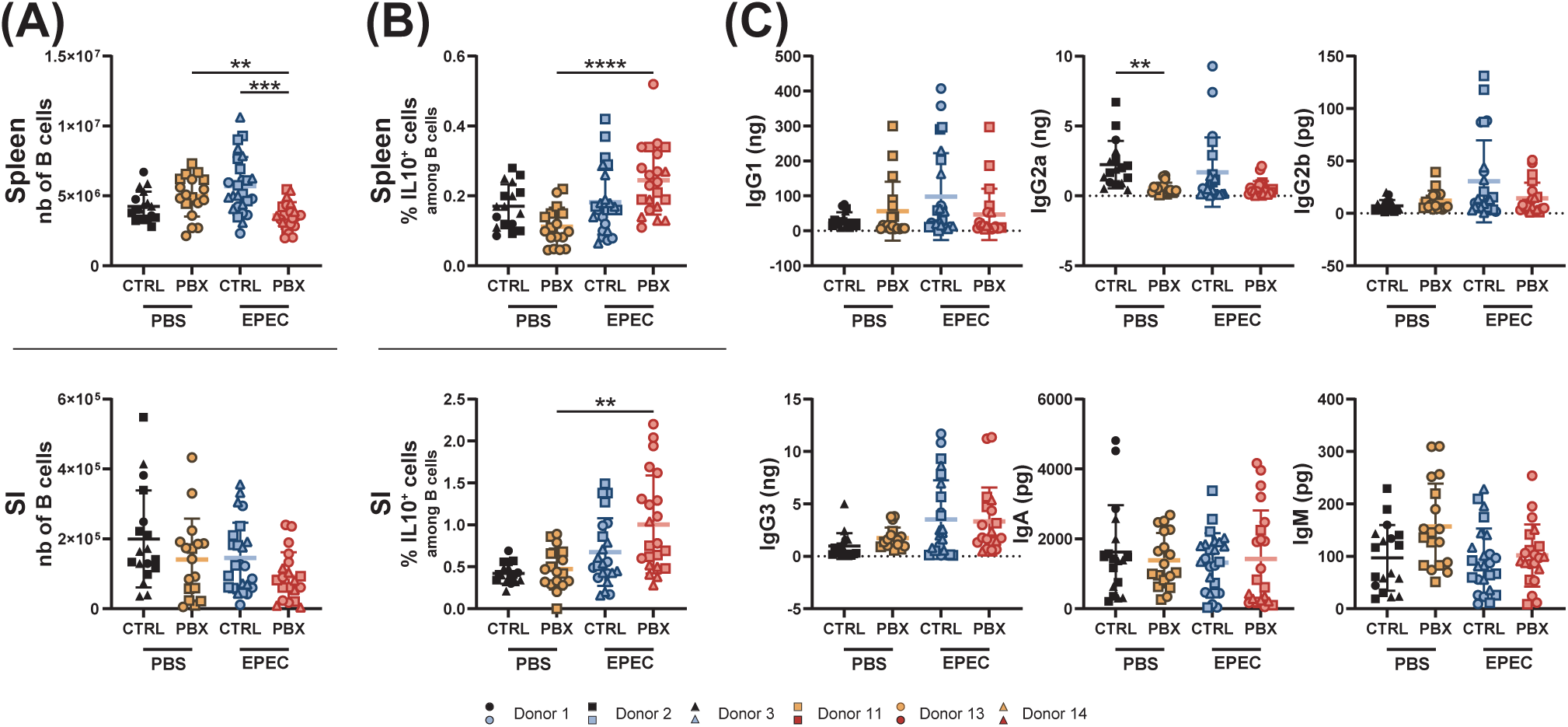
Effect of probiotic-conditioned microbiota on B cells during early life gastrointestinal infection. Spleen and small intestine (SI) of sham (PBS) or infected (EPEC) neonates at postnatal day 12 born to mothers humanized with placebo (CTRL) or probiotic (PBX) treated preterm feces were collected 7 days post-infection and analyzed by flow cytometry. **(A)** Cell counts of B cells. **(B)** Parental percentage of regulatory (IL-10^+^) B cells. **(C)** Amount of IgG1, IgG2a, IgG2b, IgG3, IgA, and IgM in the total cecal content. Results are pooled from 1 (Donors 1, 11, 14), 2 (Donors 2, 13) or 3 (Donor 3) independent experiments and are presented as mean ± SD. Each symbol represents one mouse. Sample sizes per donor are given as *n*PBS/*n*EPEC: CTRL donors - Donor 1 (●, 3/7), Donor 2 (▪, 8/9), Donor 3 (▴, 7/10); PBX donors - Donor 11 (□, 7/5), Donor 13 (○, 11/11), Donor 14 (△, 0/6) (** = p<0.005; *** = p <0.001; **** = p <0.0001).

